# On Using Transfer Learning For Plant Disease Detection

**DOI:** 10.1101/2020.05.22.110957

**Authors:** Abhinav Sagar, Dheeba Jacob

## Abstract

Deep neural networks has been highly successful in image classification problems. In this paper, we show how neural networks can be used for plant disease recognition in the context of image classification. We have used publicly available Plant Village dataset which has 38 classes of diseases. Hence, the problem that we have addressed is a multi class classification problem. We compared five different architectures including VGG16, ResNet50, InceptionV3, InceptionResNet and DenseNet169 as the backbones for our work. We found that ResNet50 achieves the best result on the test set. For evaluation, we used metrics: accuracy, precision, recall, F1 score and class wise confusion metric. Our model achieves the best of results using ResNet50 with accuracy of 0.982, precision of 0.94, recall of 0.94 and F1 score of 0.94.

## 1 Introduction

It is very important to get an accurate diagnosis of plant diseases for global health and well being. In this ever changing environment, identifying the disease including early prevention is important to avoid problems that we might face otherwise. Some of these problems could have devastating impacts on humanity including global shortage of food. It is crucial to prevent unnecessary waste of financial resources achieving a healthier lifestyle, by addressing climate change from an ecological perspective. It is difficult for the naked eye of a human being to catch all sorts of problems with plant diseases. Also doing this time and time again is also laborious and unproductive work. In order to achieve accurate plant disease detection, a plant pathologist should possess good observation skills so that one can identify characteristic symptoms. An automated system designed to help identify plant diseases by the plant’s appearance and visual symptoms could be of great help. This can be deployed in agricultural fields so that the whole pipeline can be automated. This would not only lead to better efficiency as machines could perform better than humans in these redundant tasks but also improve the productivity of the farm. Our work solves the above mentioned problem of automating plant disease classification using deep learning and computer vision techniques.

### 1.1 Plant Disease Detection

Disease detection in plants plays an important role in agriculture as farmers have often to decide whether the crop they are harvesting is good enough. It is of utmost importance to take these seriously as it can lead to serious problems in plants due to which respective product quality, quantity or productivity is affected. Plant diseases cause a periodic outbreak of diseases leading to large-scale death which severely affects the economy. These problems need to be solved at the initial stage, to save the lives and money of people. Automatic classification of plant diseases is an important research topic as it is important in monitoring large fields of crops and at a very early stage, if we can detect the symptoms of diseases when they appear on plant leaves. This enables computer vision algorithms to provide image-based automatic inspection. Comparatively, manual identification is labor intensive, less accurate and can be done only in small areas at a time. By this method, the plant diseases can be identified at the initial stage itself and the pest and infection control tools can be used to solve pest problems while minimizing risks to people and the environment.

## 2 Existing Work

A lot of research has been done in the last decade on plant disease detection using deep learning and computer vision. Machine Learning approaches include traditional computer vision algorithms like haar, hog, sift, surf, image segmentation, Support Vector Machines (SVM), using K-Nearest Neighbours (KNN), K-means and Artificial Neural Networks (ANN). Deep Learning based plant disease classification models includes the use of a variety of CNN models such as AlexNet, GoogleNet, VGGNet etc. It is seen oftentimes as the dataset size is not enough, multi class classification with a lot of classes requires careful hyperparameter tuning to avoid overfitting as the model could easily get stuck in a local minimum.

(Revathi and Hemalatha, 2012) used RGB feature extraction techniques to identify the diseases in which the captured images are processed for enhancement first. After this color image segmentation is carried out to get target regions around the original image. Next homogenization techniques including Sobel and Canny filters was applied to identify the edges. These extracted edge features are used in classification to identify the disease spots present in the image. (Phadikar and Sil, 2008) uses images of the infected rice plants by digital camera. Image segmentation techniques including K means algorithms to detect infected parts of the plants was used. Finally the infected part of the leaf has been used for the classification purpose using a deep neural network using a softmax layer. (Al-Hiary et al., 2011) starts by identifying the mostly green colored pixels. Next, these pixels are masked based on specific threshold values that are computed using Otsu’s method. Then those green pixels which are more than some threshold value are masked. An additional step is that the pixels with zeros red, green and blue values and the pixels on the boundaries of the infected cluster were completely removed. (Deshpande et al., 2014) starts by capturing images that are processed for enhancement. Then image segmentation using neural networks is carried out to get target regions (disease spots) on the leaves and fruits. Concluding, if the diseased spot on leaf is bordered by yellow margin then it is said that leaf is infected by bacterial blight otherwise not. (Gavhale et al., 2014) uses image preprocessing including RGB to differentiate color space conversion, image enhancement, segment the region of interest using K-mean clustering for statistical usage to determine the defect and severity areas of plant leaves, feature extraction and classification. Finally classification is achieved using SVM. (Reyes et al., 2015) uses a pre-trained convolutional neural network using 1.8 million images and uses a fine-tuning strategy to transfer learned recognition capabilities from general domains to the specific challenge of plant identification task.

(Sagar, 2019) is known as This work uses various transfer learning architectures and shows how hyperparameter tuning can be done to achieve the best results. We demonstrate state of the art results for this particular problem.

## 3 Proposed Method

We have used the concept of transfer learning for the classification. The main advantage in using transfer learning is that instead of starting the learning process from scratch, the model starts from patterns that have been learned when solving a different problem which is similar in nature to the one being solved. This way the model leverages previous learnings and avoids starting from scratch. In image classification, transfer learning is usually expressed through the use of pre-trained models. A pre-trained model is a model that was trained on a large benchmark dataset to solve a similar problem to the one that we want to solve. We used five pre-trained models-Inception v3, InceptionResNet v2 and ResNet50, MobileNet and DenseNet169 as the pre-trained weights for our work.

### 3.1 Dataset

A public dataset is provided which contains 54,305 images of diseased and healthy plant leaves collected under controlled conditions. The images cover 14 species of crops, including: apple, blueberry, cherry, grape, orange, peach, pepper, potato, raspberry, soy, squash, strawberry and tomato. It contains images of 17 basic diseases, 4 bacterial diseases, 2 diseases caused by mold, 2 viral diseases and 1 disease caused by a mite. Each class label is a crop-disease pair, and we make an attempt to predict the crop-disease pair given just the image of the plant leaf. Figure 1 shows all the classes present in the PlantVillage dataset.

**Figure 1:**
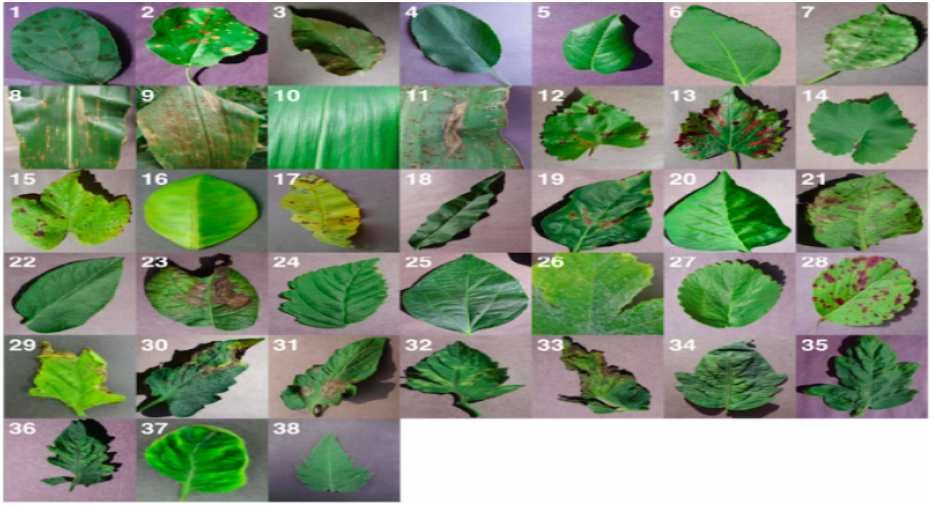
All the classes of plant disease present in dataset

### 3.2 Image augmentation techniques

The images are resized to 256 × 256 pixels, and we perform both the model optimization and predictions on these downscaled images. We used data augmentation like shearing, zooming, flipping and brightness change to increase the dataset size to almost double the original dataset size. Data augmentation techniques are often used together with traditional machine learning algorithms or deep learning algorithms to improve the accuracy of classification. In this study, the image augmentation method was used by using the Keras deep learning library in Python. Width and height change, cutting, zooming, horizontal turning, brightness, and filling operations were performed for normal class images. The image rotation degree was set to be randomly generated from 0 to 45.

In this study, the image augmentation techniques were applied only to normal images in order to balance the distribution of the samples over the classes. The number of normal samples in the dataset was increased from 1,583 to 4,266 by performing the image augmentation techniques. In this manner, the number of samples for each class was equalized. This equal distribution makes it possible to use all of the data instead of selecting random data during the training process. It is expected that this situation increases the accuracy of the training and positively affects the classification results. Image augmentation techniques are shown in Fig 2.

**Figure 2:**
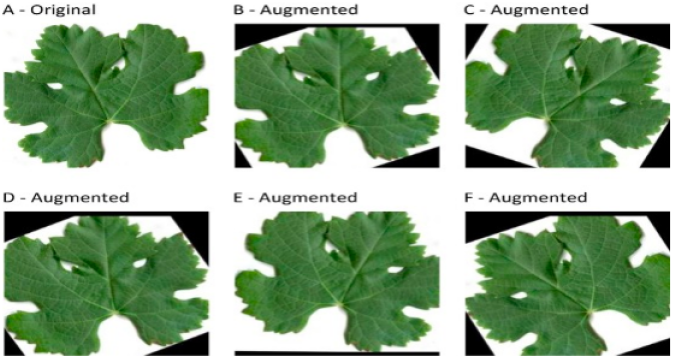
Data augmentation

### 3.3 Optimization

The main purpose of optimization methods is to update the weights at every stage until the best learning in CNN is realized. Each method performs an update process. In the Stochastic Gradient Descent (SGD) method, the weights update is performed in every iteration for each instance present in the training set. Because of this reason, it tries to achieve the goal as early as possible.

### 3.4 Dropouts

Dropout is one of the common regularization techniques to prevent neural networks from overfitting. Others often used regularization methods like L1 and L2 to reduce overfitting by penalizing the cost function. Dropout on the other hand, modifies the network itself by randomly dropping neurons from the neural network during training in each iteration. When we drop different sets of neurons, it’s equivalent to training an ensemble of neural networks and thus it helps in reducing variance. The different neural networks will overfit in different ways, so the net effect of dropout will be to reduce overfitting.

### 3.5 Visualization of Feature Maps

The feature maps help in explaining what the model is learning at every layer. As the depth increases, the model is able to learn more spatial information. In other words, the neural networks go from learning edges and blobs in the first layer to complete objects in the last layers. Visualization of feature maps is important to understand what the filters are learning at each layer. The hyperparameter tuning is easier because when an error is made by the neural network we can get the reason for going wrong. The functionality and expected behaviour of the neural networks can be explained especially to non-technical stakeholders who wouldn’t accept deep learning algorithms results until there is a reasoning behind them. This also makes extending and improving the overall design of models since we’d have knowledge of the current design, including how it performs. By visualising the learned weights we can get some idea as to how well our network has learned. For example, if we see a lot of zeros then we’ll know we have many dead filters that aren’t going much for our network, a great opportunity to do some pruning for model compression. Fig 3. shows the activation maps for the filters present in four convolution and four max pooling layers.

**Figure 3:**
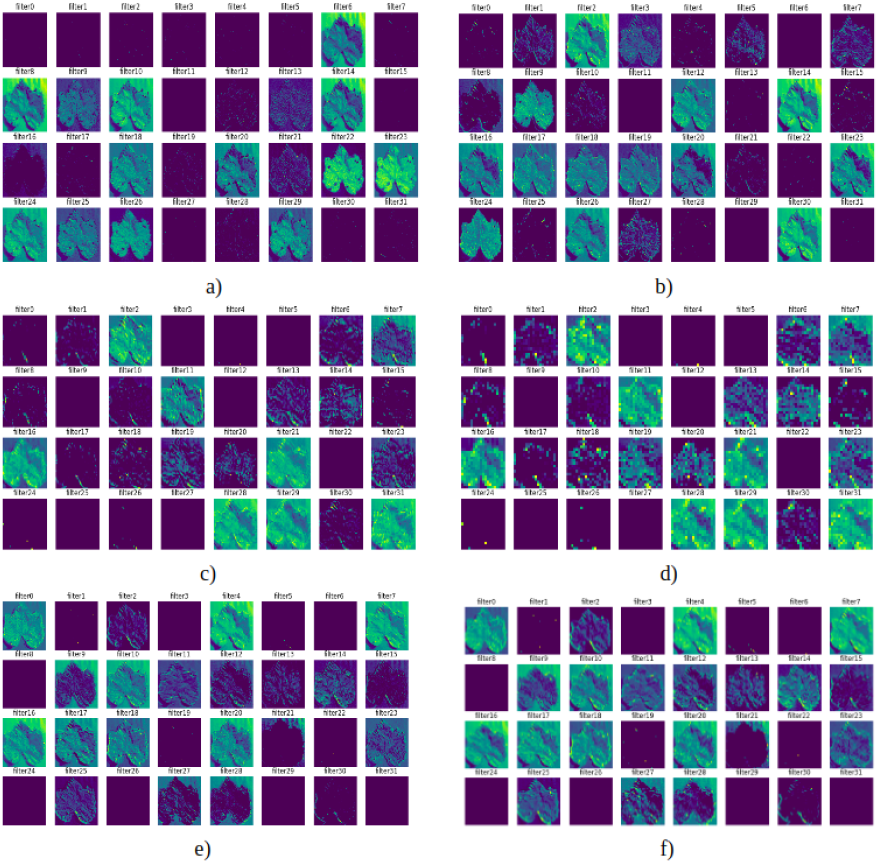
Visualization of feature maps a) First convolutional layer b) First pooling layer c) Second convolutional layer d) Second pooling layer e) Third convolutional layer f) Third pooling layer

Neural networks are often thought of as black boxes. In this way when we deploy the model in production, the feature maps come in handy as the non technical people like stakeholders, business people, doctors etc often don’t understand what neural network does behind the scenes. This makes it easy for them to get convinced and accept the results. We used two additional techniques while training ModelCheckpoint and EarlyStopping.

ModelCheckpoint is used when training requires a lot of time to achieve a good result, often many iterations are required. In this case, it is better to save a copy of the best performing model only when an epoch that improves the metrics ends. Sometimes during training we can notice that the generalization gap i.e. the difference between training and validation error starts to increase, instead of decreasing. This is a symptom of overfitting that can be solved in many ways (reducing model capacity, increasing training data, data augmentation, regularization, dropout, etc). Often a practical and efficient solution is to stop training when the generalization gap is getting worse.

### 3.6 Model Architecture

We split the data-set into three sets — train, validation and test sets. We tried with pre trained models like Inception v3, InceptionResNet v2 and ResNet 50, MobileNet and Densenet169 by fine tuning the last layers of the network. On top of the transfer learning architectures, we have added 4 custom convolutional and max pooling pooling layers. We used two dense layers with 64 neutrons and 2 neurons respectively at the last. The last layer is used for the classification with softmax as the activation function. The loss function used is binary cross-entropy. We trained the model for 20 epochs with a batch size by changing the hyper-parameters like learning rate, batch size, optimizer and pre-trained weights. We used 30 percent dropouts to reduce overfitting in between the layers and batch normalization to reduce internal covariate shift. This also helped the model avoid getting stuck in the local optimum or a saddle point. Multi class log loss was chosen as the evaluation metric. Activation function used was Relu throughout except for the last layer where it was Sigmoid as this is a binary classification problem. Further we made a couple of data generators: one for training data, and the other for testing data. A data generator is capable of loading the required amount of data (a mini batch of images) directly from the source folder, converting them into training data and training targets.

## 4 Expirimental Results

In this section we present our findings. We plotted the loss vs epochs, accuracy vs epochs and confusion matrix for the classifier. The loss vs epochs, accuracy vs epochs figure is shown in Fig 4.

**Figure 4:**
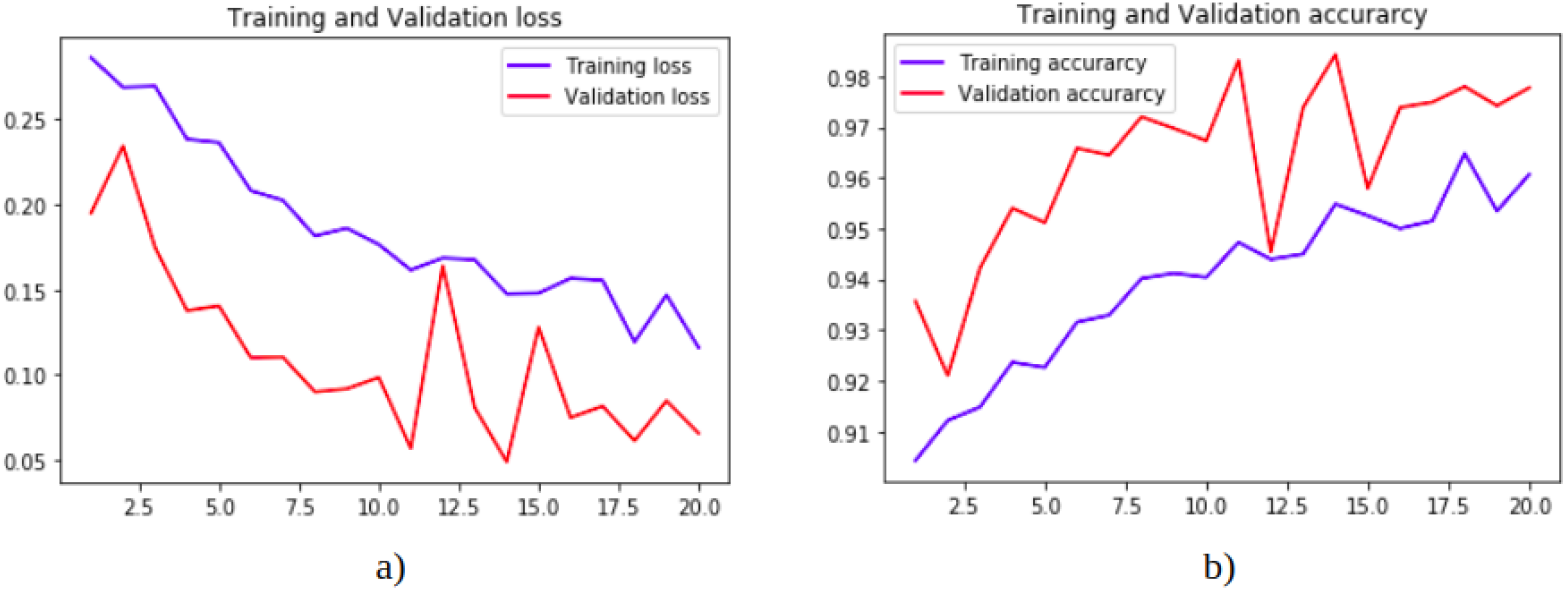
a) Loss vs epochs b) Accuracy vs epochs

We chose Multi Class Log Loss evaluation function, which is defined as:

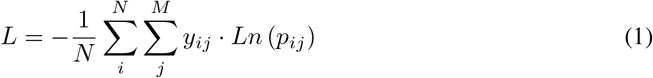

– *N* is the number of classes present in the test set
– *M* is the number of class labels which is 38 in this case
– *y*_*ij*_ depicts if the *i*^*th*^ object in the test set belongs to the *j*^*th*^ label which is given by a boolean value.
– *p*_*ij*_ is the probability that the *i*^*th*^ object belongs to the *j*^*th*^ label.
– *Ln* is the natural logarithmic function.

For a better look at misclassification, we often use the following metric to get a better idea - true positives (TP), true negatives (TN), false positive (FP) and false negative (FN). Precision is the ratio of correctly predicted positive observations to the total predicted positive observations. Recall is the ratio of correctly predicted positive observations to all the observations in actual class. F1-Score is the weighted average of Precision and Recall.

In this paper, a total of 19 types of plant disease categories are used. The evaluation and results of trained models is calculated by common classification metrics. The acronyms, TP, TN, FP, FN refer to true positive, true negative, false positive and false negative. We also used F1 for evaluation which combined both precision and recall in a single term. The higher the F1-Score, the better the model. For all three metrics, 0 means the model is performing the worst while 1 means it is performing the best. The precision, recall, F1 and support values for all the individual classes is shown in Fig 5. The average accuracy and weighted class wise accuracy is found to be 0.91 and 0.94 respectively.

**Figure 5:**
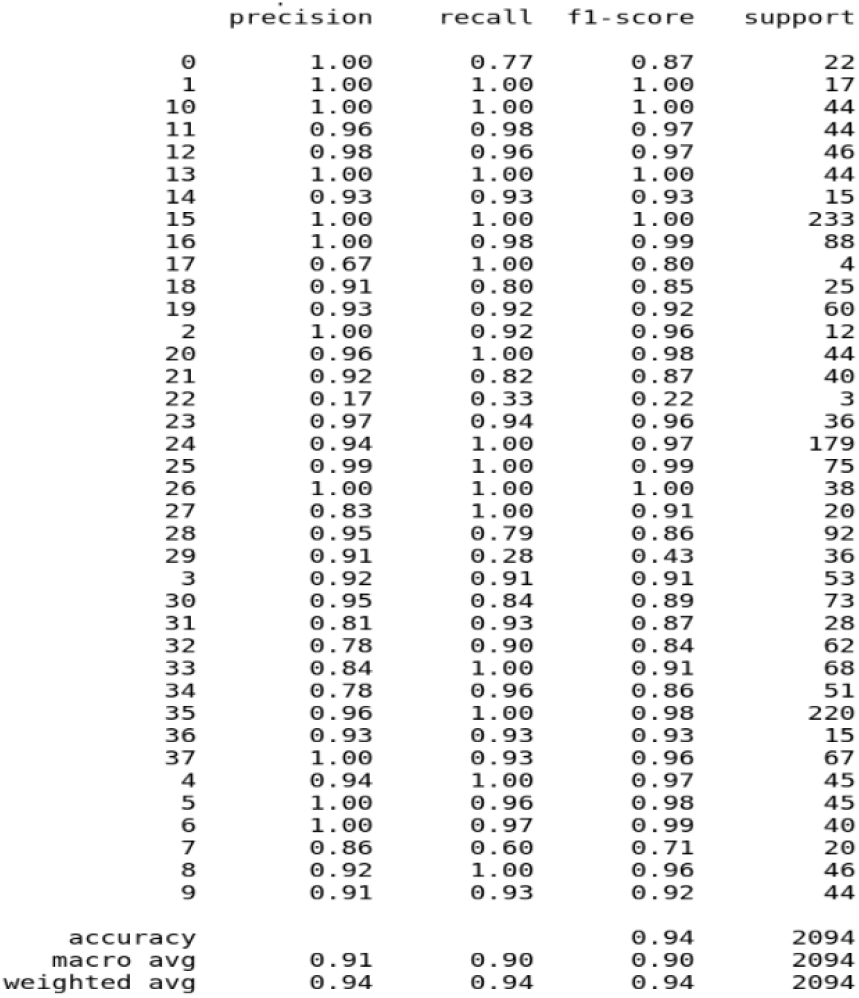
Evaluation metrics results

Next we plotted the confusion matrix for all the pairs having and not having disease. Confusion Matrix is often used as an evaluation metric when analyzing misclassification between classes. Each row of the matrix represents the examples in the class being predicted while each column represents the examples in the class which are original. The diagonals show the classes which have been classified correctly. This tells us both which classes are being misclassified and also what they are being misclassified as. The class wise confusion matrix is shown in Fig 6. Each of these contain a pair both having the disease and not having one.

**Figure 6:**
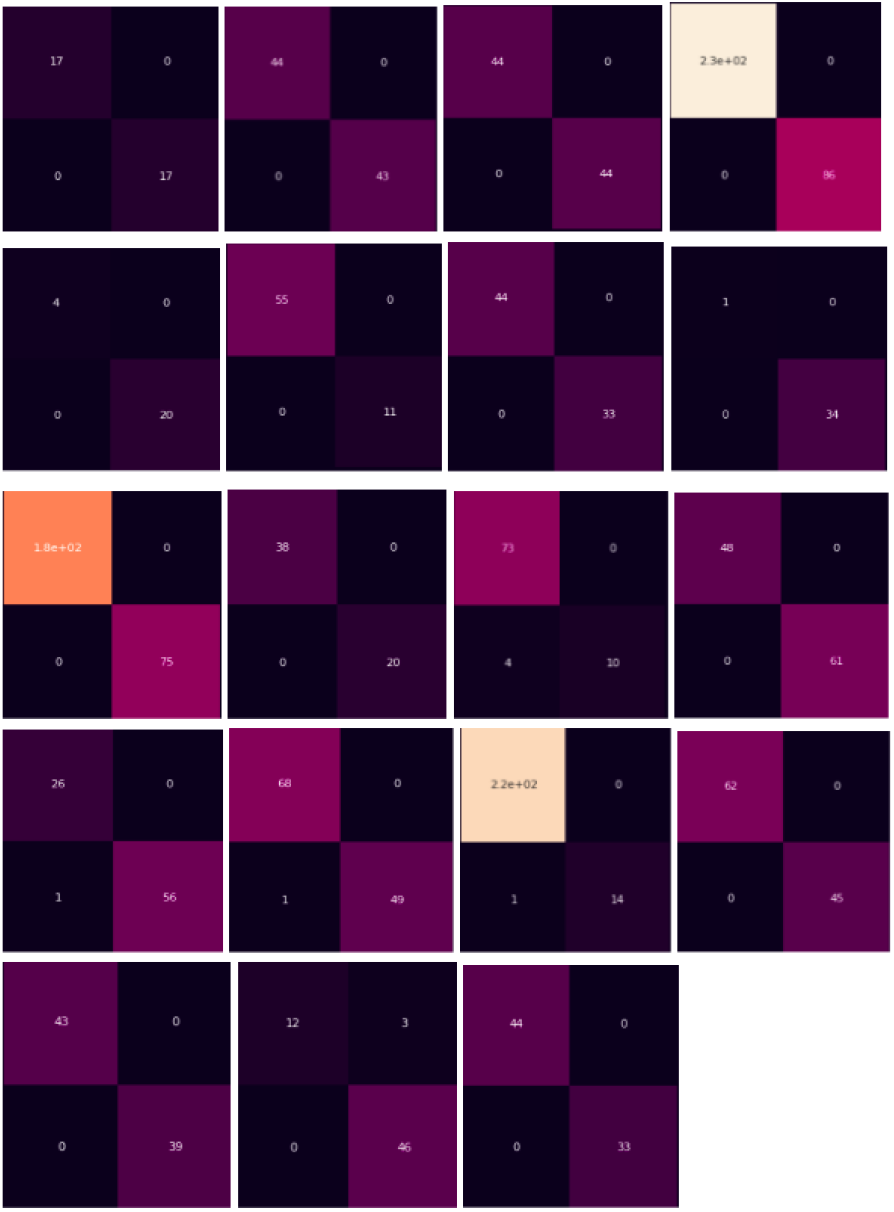
Confusion Matrix for all the classes

Finally we present our final results i.e. comparing accuracy, precision, recall and F1 score for all the transfer learning architectures used - Inception v3, InceptionResNet v2 and ResNet50, MobileNet and DenseNet169 which is shown in Table 1.

**Table 1:** Comparison of pre trained weights on results

| Evaluation Metrics | InceptionV3 | InceptionResN etV2 | ResNet50 | MobileNet | DenseNet169 |
| --- | --- | --- | --- | --- | --- |
| Accuracy | 0.971 | 0.978 | 0.982 | 0.971 | 0.974 |
| Precision | 0.92 | 0.91 | 0.94 | 0.94 | 0.92 |
| Recall | 0.94 | 0.93 | 0.94 | 0.93 | 0.93 |
| F1 | 0.93 | 0.92 | 0.94 | 0.93 | 0.93 |

## 5 Conclusions

Diseases in plants are a major threat to food supply worldwide. This paper demonstrates the technical feasibility of deep learning using convolutional neural network approach to enable automatic disease diagnosis through image classification. Using a public dataset of 54,306 images of diseased and healthy plant leaves, a deep convolutional neural network is trained to classify crop species and disease status of 38 different classes containing 14 crop species and 26 diseases, achieving an accuracy of 98.2 percent with residual network architecture. In this paper, a new approach of using deep learning methods was explored in order to automatically classify and detect plant diseases from leaf images. The developed model was able to distinguish between healthy leaves and different diseases, which can be visually diagnosed. The complete procedure was described, respectively, from collecting the images used for training and validation to image augmentation and finally the procedure of training the deep CNN and fine-tuning. We summarized the final results and came to the conclusion that ResNet50 achieves the highest accuracy as well as precision, recall and F1 score.

## Acknowledgments

We would like to thank Nvidia for providing the GPUs.

## References

A. F. Agarap. Deep learning using rectified linear units (relu). arXiv preprint arXiv:1803.08375, 2018.

H. Al-Hiary, S. Bani-Ahmad, M. Reyalat, M. Braik, and Z. Alrahamneh. Fast and accurate detection and classification of plant diseases. International Journal of Computer Applications, 17(1):31–38, 2011.

T. Deshpande, S. Sengupta, and K. Raghuvanshi. Grading & identification of disease in pomegranate leaf and fruit. International Journal of Computer Science and Information Technologies, 5(3): 4638–4645, 2014.

K. R. Gavhale, U. Gawande, and K. O. Hajari. Unhealthy region of citrus leaf detection using image processing techniques. In International Conference for Convergence for Technology-2014, pages 1–6. IEEE, 2014.

K. He, X. Zhang, S. Ren, and J. Sun. Deep residual learning for image recognition. In Proceedings of the IEEE conference on computer vision and pattern recognition, pages 770–778, 2016a.

K. He, X. Zhang, S. Ren, and J. Sun. Identity mappings in deep residual networks. In European conference on computer vision, pages 630–645. Springer, 2016b.

G. Huang, Y. Sun, Z. Liu, D. Sedra, and K. Q. Weinberger. Deep networks with stochastic depth. In European conference on computer vision, pages 646–661. Springer, 2016.

G. Huang, Z. Liu, L. Van Der Maaten, and K. Q. Weinberger. Densely connected convolutional networks. In Proceedings of the IEEE conference on computer vision and pattern recognition, pages 4700–4708, 2017.

D. P. Kingma and J. Ba. Adam: A method for stochastic optimization. arXiv preprint arXiv:1412.6980, 2014.

A. Krizhevsky, I. Sutskever, and G. E. Hinton. Imagenet classification with deep convolutional neural networks. In Advances in neural information processing systems, pages 1097–1105, 2012.

S. Phadikar and J. Sil. Rice disease identification using pattern recognition techniques. In 2008 11th International Conference on Computer and Information Technology, pages 420–423. IEEE, 2008.

P. Revathi and M. Hemalatha. Classification of cotton leaf spot diseases using image processing edge detection techniques. In 2012 International Conference on Emerging Trends in Science, Engineering and Technology (INCOSET), pages 169–173. IEEE, 2012.

P. Revathi and M. Hemalatha. Identification of cotton diseases based on cross information gain deep forward neural network classifier with pso feature selection. International Journal of Engineering and Technology, 5(6):4637–4642, 2014.

A. K. Reyes, J. C. Caicedo, and J. E. Camargo. Fine-tuning deep convolutional networks for plant recognition. CLEF (Working Notes), 1391:467–475, 2015.

A. Sagar. Icc 2019 cricket world cup prediction using machine learning, 2019.

N. Srivastava, G. Hinton, A. Krizhevsky, I. Sutskever, and R. Salakhutdinov. Dropout: a simple way to prevent neural networks from overfitting. The journal of machine learning research, 15(1): 1929–1958, 2014.

C. Szegedy, W. Liu, Y. Jia, P. Sermanet, S. Reed, D. Anguelov, D. Erhan, V. Vanhoucke, and A. Rabinovich. Going deeper with convolutions. In Proceedings of the IEEE conference on computer vision and pattern recognition, pages 1–9, 2015.

C. Szegedy, V. Vanhoucke, S. Ioffe, J. Shlens, and Z. Wojna. Rethinking the inception architecture for computer vision. In Proceedings of the IEEE conference on computer vision and pattern recognition, pages 2818–2826, 2016.

A. Veit, M. J. Wilber, and S. Belongie. Residual networks behave like ensembles of relatively shallow networks. In Advances in neural information processing systems, pages 550–558, 2016.

L. Xu, J. S. Ren, C. Liu, and J. Jia. Deep convolutional neural network for image deconvolution. In Advances in neural information processing systems, pages 1790–1798, 2014.

J. Yosinski, J. Clune, Y. Bengio, and H. Lipson. How transferable are features in deep neural networks? In Advances in neural information processing systems, pages 3320–3328, 2014.

